# Whole body transcriptomes and new insights into the biology of the tick *Ixodes ricinus*

**DOI:** 10.1101/244830

**Authors:** N. Pierre Charrier, Marjorie Couton, Maarten J. Voordouw, Olivier Rais, Axelle Durand-Hermouet, Caroline Hervet, Olivier Plantard, Claude Rispe

## Abstract

**Background:** *Ixodes ricinus* is the most important vector of tick-borne-diseases in Europe. A better knowledge of its genome and transcriptome is important for developing control strategies. Previous transcriptomic studies of *I. ricinus* have focused on gene expression during the blood meal in specific tissues. To obtain a broader picture of changes in gene expression during the blood meal, our study analysed the transcriptome at the level of the whole body for both nymphal and adult ticks. *I. ricinus* ticks from a highly inbred colony at the University of Neuchâtel were used. We also analysed previously published RNAseq studies to compare the genetic variation between three wild strains and three lab strains, including the strain from Neuchâtel.

**Results:** RNA was extracted from whole tick bodies and the cDNA was sequenced, producing 162,872,698 paired-end reads. Our reference transcriptome contained 179,682 contigs, of which 31% were annotated using Trinotate. Gene expression was compared between ticks that differed with respect to stage (nymph, adult), sex (female, male), and feeding status (unfed, partially fed). We found that blood feeding in nymphs and female adult ticks increased the expression of cuticle-associated genes. Using a set of 5,422 single nucleotide polymorphisms to calculate the heterozygosity, we found that the wild tick populations of *I. ricinus* had much higher levels of heterozygosity than the three lab populations.

**Conclusion:** Using high throughput strand-oriented sequencing for whole ticks in different stages and feeding conditions, we obtained a *de novo* assembly that significantly increased the genomic resources available for *I. ricinus*. Our study illustrates the importance of analysing the transcriptome at the level of the whole body to gain additional insights into how gene expression changes over the life cycle of an organism. Our comparison of several RNAseq datasets shows the power of transcriptomic data to accurately characterize genetic polymorphism and for comparing different populations or sources of sequencing material.

## Background

Ticks are vectors of numerous pathogenic microorganisms (*Borrelia spp., Babesia spp*., tick-borne encephalitis virus, etc.) that cause infectious diseases to both humans and animals [1]. Ticks acquire tick-borne pathogens from infected vertebrate hosts and transmit them to other animals during the blood meal [2]. During blood feeding, the tick salivary glands secrete a complex cocktail of molecules that allows them to inhibit the different components of the response of the vertebrate host including coagulation, inflammation, and immunity [3, 4]. Tick-borne pathogens bind to these tick salivary gland proteins to evade host immunity and enhance their own transmission [5, 6]. Similarly, during acquisition, tick-borne pathogens interact with tick proteins that allow them to persist in the tick midgut [7]. For this reason, anti-tick vaccines have traditionally targeted tick proteins in the salivary glands or midgut in the hopes of reducing the efficiency of tick feeding and pathogen transmission [8]. However, a broader knowledge of genes involved in other aspects of the tick life cycle (e.g. growth and moulting) may lead to alternative control strategies.

*Ixodes ricinus* is one of the most abundant and widespread tick species in Europe where it transmits a number of tick-borne diseases including Lyme borreliosis and tick-borne encephalitis [9, 10]. Hard ticks of the genus *Ixodes* have three motile stages: larva, nymph, and adult. The immature stages (larvae and nymphs) take a single blood meal, then moult to the next stage; adult females take a blood meal to produce eggs. There is currently much interest in studying the genome of *I. ricinus* and other tick species in the hopes of developing vector control strategies. Recent advances in sequencing technology have made it possible to study large catalogues of gene transcripts (the transcriptome) in individual species. By comparing gene expression between different states (e.g. developmental stage, sex, environmental conditions, etc.), these studies can provide insight into gene function. These RNA sequencing studies can also provide information on genetic variation within and among populations [11]. To date, several studies have investigated gene expression in *I. ricinus* [12, 13, 14, 15, 16, 17]. Most of these studies have focused on gene expression during the blood meal in either the nymphal tick or the adult tick, which is when pathogen transmission occurs. The majority of these studies have investigated gene expression in the tick salivary glands and/or the tick midgut because these tissues are critical for pathogen transmission [12, 13, 14, 16]. Taken together these studies have shown that there are thousands of transcripts that are differentially expressed with respect to the duration of the blood meal, the developmental stage (nymph versus adult), the specific tissue (salivary glands versus midgut), and other conditions [12, 13, 18, 16, 14, 15, 17, 19]. Most of these studies have focused on gene expression during the blood meal in either the nymphal tick or the adult tick, which is when pathogen transmission occurs. The majority of these studies have investigated gene expression in the tick salivary glands and/or the tick midgut because these tissues are critical for pathogen transmission [14].

The purpose of the present study is to explore new transcriptomic data of *I. ricinus* to improve improve several aspects of the existing knowledge. First, we wanted to enrich the global catalogue of genes for *I. ricinus*. We used whole tick bodies, which is expected to provide a broader description of the transcriptome compared to the previously published tissue-restricted libraries. We used strand-oriented sequencing, which produces contigs in the direction of transcription for the majority of transcripts. This type of sequencing gives a higher accuracy in the process of gene identification, especially for genes without detectable homology. Second, our design included different developmental stages and both sexes in order to capture the highest possible transcriptional diversity. Our study is therefore expected to give new insights into sex-specific gene expression (with the first high throughput transcriptome sequences for *I. ricinus* males and stage-specific gene expression adult females) at the level of the whole body (nymphs versus adult females). The third innovative aspect of our study was the exploration of high throughput transcriptomic sequencing to detect and compare polymorphism levels among different sources of ticks. Transcriptome sequencing can identify single nucleotide polymorphisms (SNPs) on thousands of coding genes - see [14] for an application for tick data. In the present study, we first studied polymorphism in a highly inbred laboratory strain of *I. ricinus*, which was expected to show low heterozygosity. We then compared the results from our study to the results from three previously published RNA-Seq studies that used different sources of *I. ricinus* tick material: i. wild ticks, ii. F1 offspring obtained from a mating between two wild ticks, and iii. a tick cell line. Specifically, we compared levels of polymorphism and heterozygosity between 4 different sources of *I. ricinus* tick material that were expected to differ with respect to their genetic variability. This information provides a large catalogue of polymorphic sites in expressed regions, and provides an important database for future population genetic studies.

## Methods

### Origin of ticks

The *I. ricinus* ticks used in this study came from a laboratory colony reared at the University of Neuchâtel, in Neuchâtel, Switzerland. This colony was initiated in 1978 with a small number of wild ticks collected from a natural population near Neuchâtel and has been maintained as follows. Larval and nymphal ticks are fed on laboratory mice (*Mus musculus*) and adult ticks are fed on rabbits (*Oryctolagus cuniculus*). Completion of the life cycle of *I. ricinus* (from eggs to eggs) in the laboratory takes about 1 year. Each year, there is at least one cycle of sexual reproduction with a mating population of 40 to 50 adult ticks. There has been no admixture between this tick colony and wild *I. ricinus* ticks for almost 40 years (~40 generations). The ticks from this colony are therefore expected to be pathogen-free and have reduced genetic diversity due to prolonged inbreeding.

### Ethics statement and animal experimentation permits

The use of vertebrate animals (mice and rabbits) to maintain the *I. ricinus* colony at the University of Neuchâtel was performed following the Swiss legislation on animal experimentation. The commission that is part of the ‘Service de la Consommation et des Affaires Vétérinaires (SCAV)’ of the canton of Vaud, Switzerland evaluated and approved the ethics of this part of the study. The SCAV of the canton of Neuchâtel, Switzerland issued the animal experimentation permit (NE05/2014).

### Preparation of the biological samples

We obtained ticks in five different biological states (sample size in brackets): i. unfed nymphs (n=60), ii. partially fed nymphs (n=12), iii. unfed adult females (n=8), iv. partially fed adult females (n=4), and v. unfed adult males (n=12). Samples sizes for cDNA sequencing differed among biological states because the amount of mRNA per individual differed between stages (nymphs versus adults) and feeding conditions (unfed versus partially fed). Nymphs were fed on mice (*Mus musculus*) and removed after 24 hours of attachment while adult females were fed on rabbits (*Oryctolagus cuniculus*) and removed after 48 hours of attachment. These feeding durations for both stages (adult ticks feed longer than nymphs) correspond to phase 1 or the slow phase of engorgement following the two phases defined by Lees [20]. During this phase, the nymphs and adults both reach approximately 1/4 of their final engorgement size. Unfed and partially fed ticks were flash-frozen at −80°C before DNA and RNA extractions. To allow statistical comparisons of gene expression levels among the different conditions, three biological replicates were obtained for each of the five combinations of stage and feeding status (a total of 15 samples were prepared).

### Total RNA extraction

Whole tick bodies were ground with a soft plastic pestle in Trizol (Invitrogen, Life Technologies, Carlsbard, CA, USA) on dry ice. RNA was purified as follows: after adding chloroform, the ground material was centrifuged, the aqueous phase was transferred into an RNase-free tube and was topped up with ethanol. RNA was extracted using a NucleoSpin RNA XS column (Machery-Nagel), which included a DNAse treatment. A second DNase treatment (Machery-Nagel) in RNasin (Promega) was performed to ensure complete degradation of any remaining genomic DNA. The absence of genomic DNA was confirmed by PCR tests that targeted the 18S ribosomal RNA gene of *I. ricinus*.

### Library preparation and sequencing

The quantity and quality of extracted RNA was evaluated with NanoDrop (Thermal Scientific, DE), Qubit (Invitrogen, CA) and Experion machines (Bio-RAD Laboratories Inc., USA). All samples had sufficient quantities, concentrations and qualities of RNA to proceed with library preparation. Poly-A selection was used to target mRNAs, followed by strand-specific cDNA synthesis with an insert size of 150–400 bp, PCR amplification and library purification. Individual tags used for the 15 samples allowed multiplex sequencing. Sequencing was done on one lane of an Illumina Hiseq 2500 machine (v4 chemistry).

### Quality and assembly of reads

To produce the data set, the raw paired-end reads (2×125 bp) were first cleaned. Adapters were clipped and low quality regions were filtered using Trimmomatic (release 0.36) [21]; only reads with a minimum of 36 high quality-scored contiguous bases were kept. Summary statistics of the sequence quality were checked for each library by visualizing the FastQC report (release 0.11.5) [22].

### Filtering out rRNA reads and *de novo* assembly

To filter out reads corresponding to rRNA gene expression, reads were mapped to a large contig encompassing the 18S, 28S and 5.8S ribosomal genes. This contig was obtained by performing a preliminary *de novo* assembly of publicly available Illumina transcriptomic sequence data for *I*. *ricinus* as of June 2014 (see supplementary Table S1). Reads were mapped to this contig using Bowtie 2 [23]. This contig was expected to be more effective for removing the rRNA reads than the published rRNA sequences of *I. ricinus* because the former contains complete or nearly complete sequences whereas the latter only contains partial sequences. All the reads that did not map to rRNA were assembled with Trinity (release 2014–07–17) [24], using the ‘dUTP library preparation’ option to take into account strand-oriented sequencing.

### Assessment of transcriptome completeness

A common test used to assess the “coverage” (or information completeness) of a given sequence data set is to analyse random samples of reads, and to create a saturation curve describing the relationship between different metrics (numbers of contigs of a specified length, number of matches to a known set of genes, etc.) and the read sample size. Complete data sets have saturation curves that plateau more quickly compared to incomplete data sets. To determine the coverage or completeness of our data set, we randomly sampled 1, 2, 5, 10, 20, 50, 80, 100, 140, and 160 million reads from our cleaned libraries, as detailed below. Completeness was also estimated by determining the presence of homologs of conserved arthropod genes using the BUSCO approach [25]. BUSCO v1 uses a reference database of 2,675 conserved arthropod genes (BUSCO genes) and searches for potential homologs in the database of interest by running BLAST [26] and HMMER [27]. Conserved BUSCO genes are assigned to four classes of genes: i. missing, ii. fragmented, iii. duplicated, and iv. complete. To determine if the open reading frames (ORFs) were correctly predicted, we checked the strands of the predicted genes within the contigs matching the BUSCOs. To produce the final assembly, we reduced the potential redundancy resulting from the presence of alternative transcripts in the contigs. We clustered similar sequences using cd-hit-est [28], retaining the longest transcript of each cluster. Identity parameters were chosen to cluster nearly identical sequences resulting from alternative splicing. We used relatively stringent parameters for clustering: the local alignment had to comprise more than 50% of the longest alignment and more than 80% of the shortest alignment, with a minimum nucleotide identity of 98%. To assess the loss of information produced by the clustering, we checked read recruitment in our final set of contigs by mapping with Bowtie 2 [23].

### Comparison of the completeness of our assembly relative to other assembled transcriptomes

In recent years, several research groups have produced RNAseq data sets for *I. ricinus*. For most of these projects, the *de novo* assemblies (or sets of predicted genes derived from these assemblies) have been published in the Transcriptomes Shotgun Assembly (TSA) division of Genbank. Using statistics provided by BUSCO [25], we compared the completeness of our final assembly to that of six different assemblies/gene sets, which were obtained from six different RNAseq projects (see supplementary Table S4; TSA accessions: GADI01, GANP01, GBIH01, GCJ001, GEFM01 and GEGO01).

### Gene prediction and annotation

The prediction of coding sequences (>100 amino acids) was performed using TransDecoder, which is part of the Trinity software [24]. The TransDecoder options were set to account for the strand orientation of the sequencing (i.e. ORFs were searched only on the forward strand). Finally, annotations from comparison with public databases were used to filter multiple ORF predictions by transcripts (see below). Following the pipeline recommendation of Trinotate (release 2.0.2) [29], both contigs and predicted peptides were compared by blastx+ and blastp+ (release 2.2.29) [26] to releases of Swissprot and Uniref90 (available on https://data.broadinstitute.org/Trinity/Trinotate v2.0 resources/). Protein domains were identified using HMMER (release 3.0 from March 2010) [27] with PFAM-A [30], signal peptides with SignalP [31], and transmembrane domains with TMHMM [32]. We tagged ribosomal RNAs using RNAmmer [33]. All these layers of annotation were combined by Trinotate to assign gene ontology (GO) information to each contig. In the case of multiple ORF predictions for a contig, if one ORF was similar to a known protein while the others was not, only the former ORF was retained. In other cases (several ORFs with similarity to known proteins, or several ORFs with no similarity to known proteins), the different ORFs were retained. As we were particularly interested in cuticular proteins, all peptides that contained the chitin-binding domain (PF00379) were classified using the CutProtFam-Pred webserver [34].

### Differential expression and GO enrichment

In addition to the reconstruction of transcript sequences, RNA sequencing also allows the user to quantify transcript expression by counting the number of sequenced reads that map to a given transcript. Paired reads for each library were pseudo-aligned on the Transcriptome de Bruijn Graph (T-DBG) using Kallisto [35]. We chose this method, based on a k-mer approach, because it is much faster while providing the same accuracy as the best mapping approaches [35]. This method produced raw counts and normalized count statistics (TPM, or transcripts per million reads) for each assembled contig. These counts allowed us to test for differential expression between conditions. We compared the following conditions: partially fed ticks versus unfed ticks (to assess the feeding-related changes in gene expression), nymphs versus adults (to assess stage-related changes in gene expression), and unfed adult females versus unfed adult males (to asses sex-related changes in gene expression). The sex of the nymphs cannot be determined because sexual dimorphism is only present in adult ticks. Differential expression analyses were performed with the R package DESeq2 [36] using the raw counts from Kallisto. To describe the relevant biological changes between conditions, we used predictions produced by gene ontology (GO) term annotation, which included: “molecular function”, “biological process” and “cellular localization” [37]. GO terms were compared between transcripts that were not differentially expressed between conditions (unbiased transcripts) versus transcripts unbiased transcripts versus transcripts that were differentially expressed between conditions. We defined unbiased transcripts as contigs with no significant change in expression between conditions (fold change less than 2 and adjusted p-value higher than 0.05). Enrichment analysis was performed using the elim method using Kolmogorov-Smirnov tests developed and implemented by Adrian Alexa in the R package TopGO [38]. As suggested by this author, multiple testing was taken into consideration by using the false discovery rate (FDR) on the enrichment test p-values. The resulting GO enrichments were analyzed using the R package GOprofiles [39], which provides visualization tools. GO enrichment comparisons are by definition limited to contigs with assigned GOs, whereas many more contigs can show significant changes in expression between conditions. A substantial number of contigs had no assigned GOs but did have other annotations, such as domains identified through the PFAM analysis. We therefore used a text mining analysis of the PFAM domains to further compare changes in gene expression between the different conditions (unfed/fed, nymph/adult, male/female). This approach allowed us to distinguish the most common terms associated within a text. We therefore extracted the PFAM terms associated with contigs over-expressed in each of the conditions defined above (fed/unfed, adult/nymph, male/female), and treated them as a single text for each category. To prevent over-representation of transcripts with many PFAM domains, only the first ten PFAM terms were retained for a transcript. Over-expressed contigs were defined as contigs with a fold change larger than 4 and a significant p-value (p<0.05) in the DESeq2 analysis. PFAM descriptions were edited to remove the less informative terms (e.g. “protein”, “domain”, “motif’). We then used an in-house R script to draw clouds of words with word sizes proportional to their frequency in the text. This approach is complementary to the GO-enrichment tests, as it helps to visualize major shifts in expression among conditions.

### Summary of results by peptide predictions and by contigs

An annotation file with tab-separated values was produced, providing all the information from the Trinotate report, raw counts from Kallisto, log fold changes and p-values for differential expression, and BUSCO information. This report contains one line per peptide prediction and was deposited with a DOI on the Zenodo platform (see the section “Accessibility of Data” at the end of the article).

### Polymorphisms

Polymorphism was surveyed in the reads produced through our project (corresponding to an inbred line) but also for data from five other RNAseq projects of *I. ricinus* publically available in Genbank. Three published data sets used wild tick populations from Sénart in France (SEN) [40], from the Czech Republic (CZ-W) [13], and a mixture of wild tick populations (LUX) that was provided by Charles River Laboratory [17]. The other two data sets were based on F1 full sibs from a cross between wild ticks from the Czech Republic (CZ-F1), and a tick cell line (CL). More details on those 5 data sets are given in Table S3 in the supplementary material. Reads sequences are available at the NCBI Sequence Reads Archive (SRA) and organized by Bio-Project. After downloading reads from the SRA archives, the reads were cleaned using Trimmomatic (with the same parameters as above). As estimates of polymorphism and heterozygosity depend on sample size, we standardized the sample size for each data set by randomly sampling 30 millions reads from each SRA. The combined data sets were analysed to detect single nucleotide polymorphisms (SNPs). SNPs were predicted with a “direct from the reads” approach, using KisSplice (release 2.4.0) [41], a software that identifies variations by detecting “bubbles” in the De Bruijn graph. SNPs were mapped on our final set of contigs using Blat (version 36) [42]. A report assessing various parameters for each SNP (location, reliability, etc.) was provided by Kiss2refTranscriptome [43]. To minimize false positives, we retained only SNPs that respected the following criteria: i. SNPs had to be covered by at least ten reads in each data set and ii. SNPs needed to be uniquely mapped (e.g. mapping to a single component). Using this restricted set of SNPs, we used variant counts produced by KisSplice to calculate allele frequencies at each polymorphic sites, for each of the 6 RNAseq data sets. We estimated heterozygosity (using the formula He = 2pq, where p and q are the frequency of each variant) for each SNP position, and for each data set. Genetic differentiation between data sets was estimated using *F_ST_* calculated by BayesScan [44] with default parameters.

## Results

### Reads and assembly statistics

We obtained a total of 210,229,106 paired strand-oriented reads. After filtering out the poor quality reads and orphan reads, (13,611,318 reads) and after excluding the reads assigned to rRNA (33,745,090 reads), the final set contained 162,872,698 trimmed, good quality reads (see Table 1). For the fifteen libraries, the mean number of reads was 10,858,180 reads per library (range = 7,628,548 to 15,814,366 reads). Trinity produced an initial assembly set of 427,491 contigs, of which half had a length between 200 and 300 bp. As these small contigs are expected to be mostly represent gene fragments or untranslated region (UTR) sequences, we discarded them and only considered contigs above 300 bp. After reduction of redundancy step [28], the resulting final assembly contained a total of 192,050 contigs. For the 15 different tick libraries, 88% and 91% of the reads mapped back to contigs (see Table 1). This good recruitment rate suggests that i. the Trinity-produced assembly managed to capture most of the information contained by the reads, and ii. little information was lost after eliminating the smaller class of contigs (¡ 300 bp). As an internal test of completeness, we analyzed the numbers of contigs of different sizes that resulted from the assemblies of random subsets of reads of increasing sample size. There was no clear saturation when considering all contigs, as the number of contigs was still rising (with only a moderate decrease of the slope) for even the largest read sample sizes (see supplementary Figure S2). However, when only considering contigs above a certain size, the number of contigs clearly tended to plateau; this plateau can already be seen for contig size >300 bp, whereas the plateau was marked for contigs >1,000 bp. This saturation effect was also shown with the BUSCO approach where the numbers of conserved arthropod genes plateaued at a sample size of 100 million reads (see supplementary Figure S3). Overall, this result suggests that our complete set of reads tended to saturate the information on the mid-size to large transcripts, indicating that the coverage of our *I. ricinus* transcriptome was good (when considering all together the 5 combinations of stage, sex, and feeding conditions in our study).

**Table 1:** Description of the 15 libraries. The 15 libraries refer to the 5 combinations of stage, sex, and feeding condition for *I. ricinus* ticks, each replicated 3 times. The different columns refer to: library identification code, stage (nymph or adult), sex (male or female or unknown; sex is unknown for nymphs), feeding condition (unfed or partially fed), number of cleaned quality reads (Reads), percentage of reads that mapped back to the final set of transcripts with Bowtie2 (Recruitment) and number of counting events observed by Kallisto divided by the number of paired reads (Kallisto).

| Library | Stage | Sex | Condition | Reads | Recruitment<br>(%reads) | Kallisto<br>(%reads) |
| --- | --- | --- | --- | --- | --- | --- |
| A | Nymph | Unknown | Unfed | 13,839,882 | 88.1 | 86.5 |
| B | Nymph | Unknown | Unfed | 9,158,226 | 86.9 | 85.7 |
| C | Nymph | Unknown | Unfed | 13,233,880 | 88.4 | 86.5 |
| D | Nymph | Unknown | Fed | 12,665,880 | 89.5 | 87.8 |
| E | Nymph | Unknown | Fed | 8,230,200 | 91.4 | 89.9 |
| F | Nymph | Unknown | Fed | 7,762,182 | 89.3 | 88.0 |
| G | Adult | Male | Unfed | 8,191,900 | 91.5 | 89.7 |
| H | Adult | Male | Unfed | 9,360,492 | 90.7 | 89.0 |
| I | Adult | Male | Unfed | 11,228,194 | 91.6 | 90.7 |
| J | Adult | Female | Unfed | 11,604,308 | 91.6 | 89.6 |
| K | Adult | Female | Unfed | 15,814,366 | 91.2 | 88.8 |
| L | Adult | Female | Unfed | 10,277,418 | 90.9 | 88.7 |
| M | Adult | Female | Fed | 12,813,758 | 90.5 | 89.1 |
| N | Adult | Female | Fed | 11,063,464 | 94.7 | 94.5 |
| O | Adult | Female | Fed | 7,628,548 | 90.6 | 90.2 |
| Total |  |  |  | 162,872,698 | 90.4 | 88.9 |

### Taxonomic assignation

We used taxonomic information from the code name of the best hit on the Uniref90 proteins cluster. We found that 6.4% of assembled transcripts (n = 12,368) were assigned to fungi (considering a minimum of 50% of protein identity and an E-value lower than 10e-5). Fungal contamination of individual ticks has been observed in the *I. ricinus* colony at the University of Neuchâtel. To facilitate moulting, blood-engorged larvae are placed in tubes with moistened filter paper, which also facilitates the growth of opportunistic fungi, which are the most likely source of the contamination observed in the present study. We counted 887,767 reads that mapped to the fungi-like contigs, which represents only 1.2% of the total number of counting events by Kallisto, suggesting that contamination was modest. Statistics on the abundance of these fungi-like transcripts for each library are presented in the supplementary materials. After removing the 12,368 fungi-like contigs, the final assembly contained 179,682 contigs (see Table 2). A taxonomic assignation was obtained for 58,681 contigs; of these, 36.7% and 33.3% were assigned to *Ixodes scapularis* and *I. ricinus*, respectively (see Figure 1). Another 17% of the contigs had matches to “Acari” (i.e. tick and mite species other than *I. scapularis* and *I. ricinus*) and 2.6% had matches to “Insecta” (n=1,525 contigs). The insect species with the most abundant hits were: pea aphid *Acyrthosiphon pisum* (n=623 contigs), termite *Zootermopsis* (n=153), clonal raider ant *Ooceraea biroi* (n=72), Asian long-horned beetle *Anoplophora glabripennis* (n=70), and kissing bug *Rhodnius prolixus* (n=67). As expected, the mean identity of the hits reflected phylogenetic distance so that identity was highest for hits corresponding to *Ixodes* species, and lower for distant taxonomic groups. However, this expected pattern was reversed at the finest taxonomic level (genus *Ixodes*), where the mean distance to *I. scapularis* hits was lower than the mean distance to *I. ricinus* hits. We suggest that the relatively limited numbers of entries in Genbank for *I. ricinus* are causing this counter-intuitive result. We explored the distribution of % proteic identities (see supplementary Figure S5) for best hits to *I. ricinus* or *I. scapularis*. For both species, the highest peak of genes had a very high percentage of proteic identity (>95%). This peak probably corresponds to the orthologous genes found in both *I. scapularis* and *I. ricinus*. However, the distribution of proteic identities for *I. ricinus* shows a secondary peak of hits with a low proteic identity (~45%). These low identity hits cannot correspond to the same gene, but probably represent distant paralogs of genes, or genes having a similar protein domain. The presence of this secondary peak decreases the mean identity of hits to *I. ricinus* and explains why the mean identity of our data set is lower for *I. ricinus* than *I. scapularis*.

**Figure 1:**
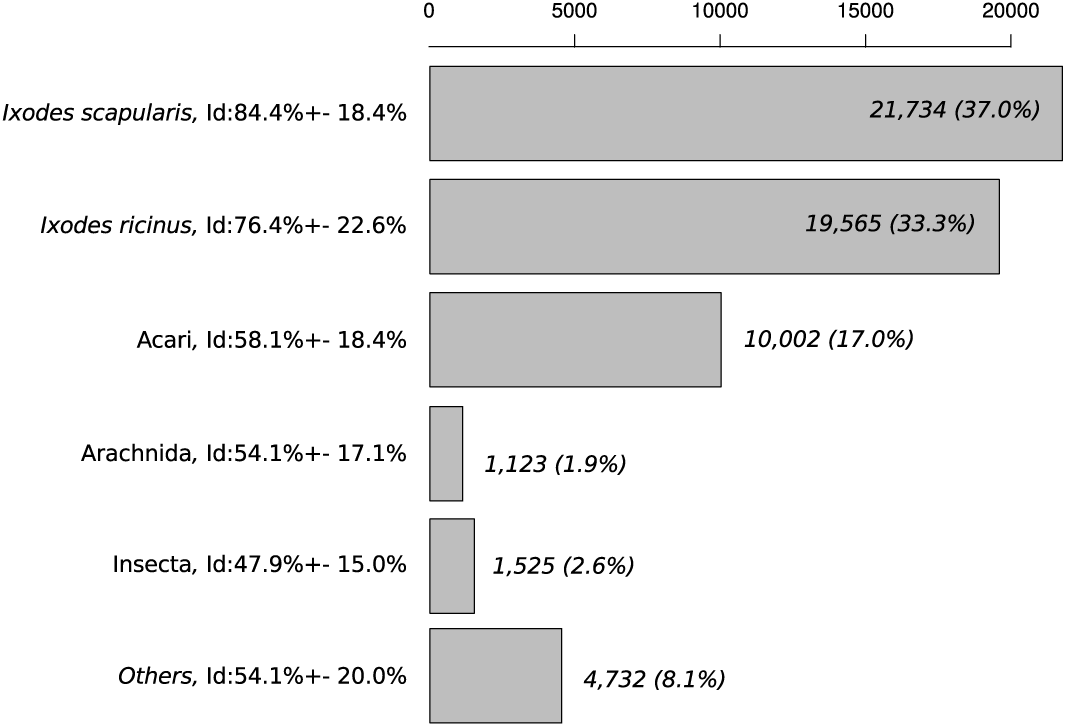
Taxonomic assignment of contigs using Uniref90 best blast hits. The bars in the bar graph correspond to the number of contigs assigned to different taxonomic groups (the percentage of contigs assigned to each taxonomic group is shown in brackets). Best hits to “Acari”, which include tick and mites are hits to Acari species other than *I. ricinus* and *I. scapularis*. Best hits to “Arachnida”, which include spiders and scorpions, are hits to Arachnida species other than Acari. The mean percentage of proteic identity and the standard deviation are shown after the taxonomic name.

### Annotation

The partially fed nymphs and adults contained blood (and therefore RNA) from mice and rabbits, respectively. The 366 contigs with high identity to mouse or rabbit proteins were removed so that the final published assembly contained 179,316 contigs. Overall, 57,098 contigs (31.8% of the total) had a significant similarity with known proteins present in Uniref90 or Swissprot (see Table 3). The percentage of contigs with a match strongly increased with contig size (Table 3). For example, the percentage of contigs with a match was as high as 68.7% for contigs longer than 1 Kb. TransDecoder predicted 57,435 peptides, of which 35.6% were predicted as complete. In total, 13,308 GO terms were extracted from 26,933 contigs (15% of all contigs). Transmembrane domains and peptide signals were found for 8,869 peptides and 3,485 peptides, respectively. Of the 179,316 contigs in the final assembly, a GO match was found for only 26,933 contigs (15.0%). In addiditon, 22,219 contigs (12.0%) had a detected PFAM domain. Some contigs had only a GO match or only a PFAM assignation, so these informations were complementary.

**Table 2:** Assembly statistics. Statistics of assembly for the final set of contigs: total size of contigs, shortest and largest contigs, mean and median contig size and the N50 contig length are expressed in base pairs.

|  | Without Fungi |
| --- | --- |
| Number of contigs | 179,682 |
| Total size of contigs | 131,092,593 bp |
| Shortest contigs | 301 bp |
| Longest contigs | 20,233 bp |
| Number of contigs > 0.55 kbp | 79,334 |
| Number of contigs > 1 kbp | 29,927 |
| Number of contig > 10 kbp | 19 |
| Mean contig size | 730 bp |
| Median contig size | 461 bp |
| N50 contig length | 873 bp |

**Table 3:** Summary of annotation statistics for the final set of contigs. For each class of contig length (class) the following data are given: number of contigs, number of contigs with significant hit on Uniref90, number of contigs with significant hit on Swissprot, number of peptides predicted by TransDecoder (Npep), and number of contigs for which gene ontology terms could be extracted (GO).

| class | contigs | Uniref90 | Swissprot | Npep | GO |
| --- | --- | --- | --- | --- | --- |
| [301, 500] | 100,348 | 19,198 (19.13%) | 5,795 (5.77%) | 16,418 | 5,412 (5.39%) |
| [501, 1000] | 49,407 | 16,666 (33.73%) | 7,033 (14.23%) | 16,335 | 6,808 (13.78%) |
| [1001, 1500] | 13,223 | 7,662 (57.94%) | 4,598 (34.77%) | 8,411 | 4,448 (33.64%) |
| [1501, 2000] | 6,448 | 4,641 (71.98%) | 3,203 (49.67%) | 5,368 | 3,186 (49.41%) |
| [2001, 2500] | 3,714 | 3,035 (81.72%) | 2,314 (62.30%) | 3,558 | 2,272 (61.17%) |
| [2501, 3000] | 2,292 | 1,933 (84.34%) | 1,533 (66.88%) | 2,388 | 1,503 (65.58%) |
| [3001, 4000] | 2,394 | 2,181 (91.10%) | 1,812 (75.69%) | 2,737 | 1,779 (74.31%) |
| [4001, 5000] | 1,025 | 956 (93.27%) | 832 (81.17%) | 1,179 | 814 (79.41%) |
| [5001, 7500] | 716 | 676 (94.41%) | 610 (85.20%) | 882 | 602 (84.08%) |
| [7501, 10000] | 96 | 93 (96.88%) | 91 (94.79%) | 132 | 91 (94.79%) |
| [10001, 25000] | 19 | 19 (100.00%) | 17 (89.47%) | 27 | 18 (94.74%) |
| Total | 179,682 | 57,060 (31.76%) | 27,838 (15.49%) | 57,435 | 26,933 (14.99%) |

### Completeness

Of the 2,675 conserved arthropod genes in the BUSCO database, our assembly contained 2,033 complete genes (completeness of 76%) and 250 partial genes (extended completeness of 85.3%). Previously published assemblies or collections of predicted genes (TSA archives) for *I. ricinus* all produced lower percentages of complete conserved genes (Figure 2). A comparison of the overlap between our new data set and the combined TSA data sets found that our assembly contains 295 BUSCO genes (11%) not present in any of these TSA data sets, whereas the combined TSA data sets contained 231 BUSCO genes not present in our assembly. We obtained a similar percentage of BUSCO contigs with correctly predicted strands (98.7%) compared to the combined TSA data sets (mean of 98.6%, minimum of 95.8% for GEGO, maximum of 100% for GANP).

**Figure 2:**
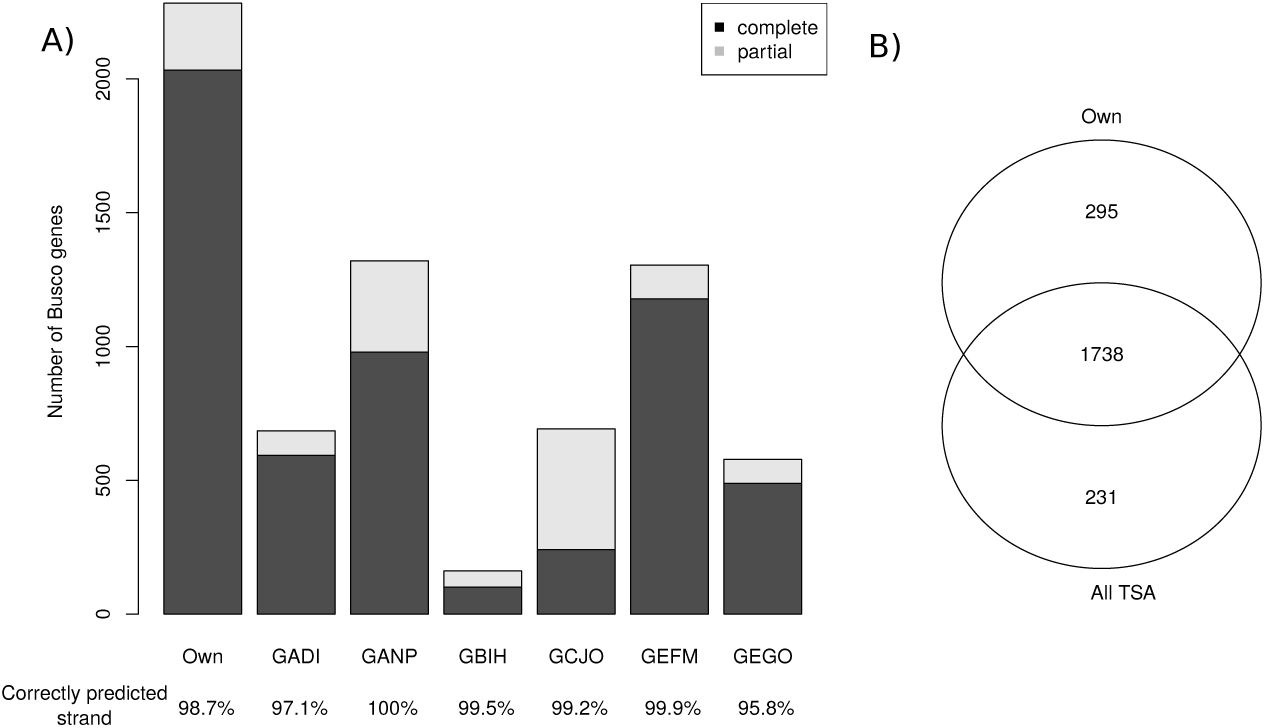
Completeness of our assembly of the *I. ricinus* transcriptome compared to the publicly available Transcriptome Shotgun Assembly (TSA) sequence database. Completeness was assessed by comparison to the to the BUSCO database, which contains 2,675 conserved arthropod genes. A) Number of BUSCO genes found in our assembly (“Own”) and the six TSA projects. The bottom of each bar shows the percentage of complete BUSCO genes predicted on the correct strand. B) Venn diagram showing the overlap between the number of complete BUSCO genes found in our assembly and in the combined TSA assemblies Altogether, the 7 transcriptome assemblies contained 2,264 BUSCO genes (completeness of 84.6%).

### Differential expression

When comparing the 200 most expressed transcripts, the “I” library of unfed adult males showed a strong deviation from all other libraries, including the two other replicate libraries of unfed adult males (libraries “G” and “H”) (see supplementary Figure S8). The “I” library also contained a particularly high percentage of rRNA reads (>60%) suggesting that this sample was of lower quality (see supplementary Figure S1). For this reason, we decided to exclude the “I” library from the analyses of differential gene expression. Comparing the levels of gene expression among libraries, we found that libraries clustered by condition for unfed nymphs, fed nymphs, and unfed females (supplementary Figure S9). Such clustering was not observed for the unfed males, whereas only two of the three samples clustered together for the fed females. The numbers of unbiased transcripts for the different comparisons were as follows: 39,719 for unfed versus partially fed, 24,801 for males versus females, and 43,992 for nymphs versus adults. We found that 11,322 transcripts (6.3%) were differentially up- or down-expressed between the unfed condition and the partially fed condition (adjusted p-value = 0.05). The numbers of differentially expressed (DE) transcripts were lower in the other two comparisons: 5,957 DE transcripts between males and females (3.3%), 2,291 transcripts between nymphs and adults (1.3%). Using the GO terms associated with DE transcripts between unfed and partially fed ticks, we found a significant enrichment (FDR < 0.05) for 2 molecular function (MF) terms, 59 biological process (BP) terms, and 4 cellular component (CC) terms (detailed results in Tables S5 to S22 in the supplementary materials). We presented a detailed view of significantly enriched GO terms (using up and down DE transcripts) between partially fed and unfed ticks without distinction of stage and sex (see Figure 3). For this analysis, the categories of the GO terms included biological process, molecular function, and cellular component, for which the false discovery rates were < 0.01, < 0.05, and < 0.05, respectively. The comparison between unfed and partially fed ticks found a significant enrichment of terms associated with cuticle production (Figure 3); specifically, the expression of cuticle-associated genes was significantly higher in partially fed ticks than unfed ticks.

**Figure 3:**
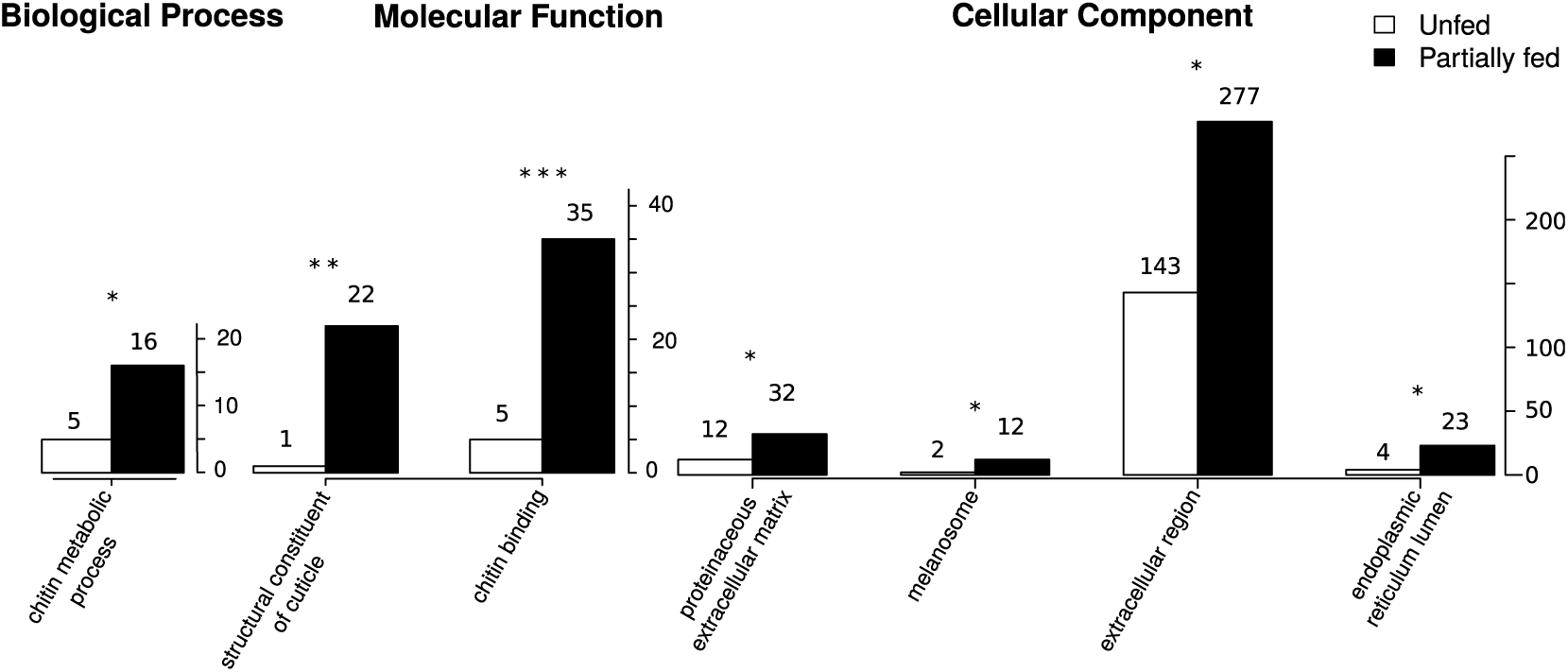
GO enrichment for the comparison between unfed and partially fed ticks. The X-axis shows the categories of the GO terms, which include: “Biological Process”, “Molecular Function”, and “Cellular Component”. The Y-axis shows the number of GO terms found in differentially expressed transcripts for unfed and partially fed ticks. The false discovery rate (FDR) was < 0.01 for “Biological Process”, < 0.05 for “Molecular Function”, <0.05 for “Cellular Component”. We represented only the best FDR for “Biological Process” (those with a corrected p-value less than 0.01 instead of 0.05). Indeed, many GO terms were obtained and did not help to visualize the result (see Table S6 and S9 in Supplementary Materials for detailed GO results). FDR significance: *<0.05; **<0.0005, ***<0.000005.

Out of 39 transcripts containing a chitin-binding PFAM domain (PF00379), one contained two PF00379 domains and 22 contained a signal peptide. All these transcripts were classified as members of group 2 using the CutProFam-Pred webserver. Expression levels in each library of cuticle-related transcripts were shown in a heatmap (Figure 4). Of the three most expressed cuticular transcripts, two transcripts shared a high identity with the proteins described by Andersen and Roepstorff [45]. The c252234_g1_i1 transcript in our study had 97.03% identity with the Ir-ACP10.9 protein (belonging to the CU109 cluster of Uniprot) [45]. This transcript was found to be highly over-expressed in partially fed ticks (588-fold change between unfed and partially fed ticks; p-value of 1.25e-43). Another transcript (c295844_g3_i4) that was highly over-expressed in partially fed adult females had 99.35% of identity with the Ir-ACP16.8 protein (belonging to the CU168 cluster of Uniprot) described in the same study [45]. Text mining of the identified domains in the differentially expressed transcripts provided complementary information (see Figure 5 and Figure S10-S14 in the supplementary material). For the genes that were over-expressed in partially fed ticks compared to unfed ticks (Figure 5), we highlight the abundance of the following terms: reprolysin and metallopeptidase and Kunitz-BPTI (these three terms were often associated in the same transcripts)as well as tick-histamine-binding, immunoglobulin, and chitin. For the genes over-expressed in unfed ticks (Figure S10), the most common terms were immnoglobulin, zinc-finger and leucine-rich-repeat. Actually, some of the most highly DE genes in unfed ticks had several IgC domains as is the case for a homolog of the gene Turtle. However, this trend was not caused by just one or a few transcripts, but by several genes with a similar structure (probably a gene family). Other comparisons of GO terms in over-expressed transcripts (nymph/adult, female/male) showed no statistical differences after correction for multiple testing (Tables S5 to S22). Text-mining of the associated domains gave additional insights into the associated gene functions. For genes that were over-expressed in nymphs (Figure S11), adult ticks (Figure S12) and unfed female (Figure S13), the zinc-finger term was by far the most abundant. Finally, domains of the transcripts over-expressed in males (Figure S14) had a high frequency of the trypsin term. Most of these transcripts matched the trypsin domain PF00089.21, whereas additional transcripts matched the domain BP-Kunitz-PTI (trypsin inhibitor) (PF00014.18). Thus, different types of protein contributed to the high frequency of the term trypsin.

**Figure 4:**
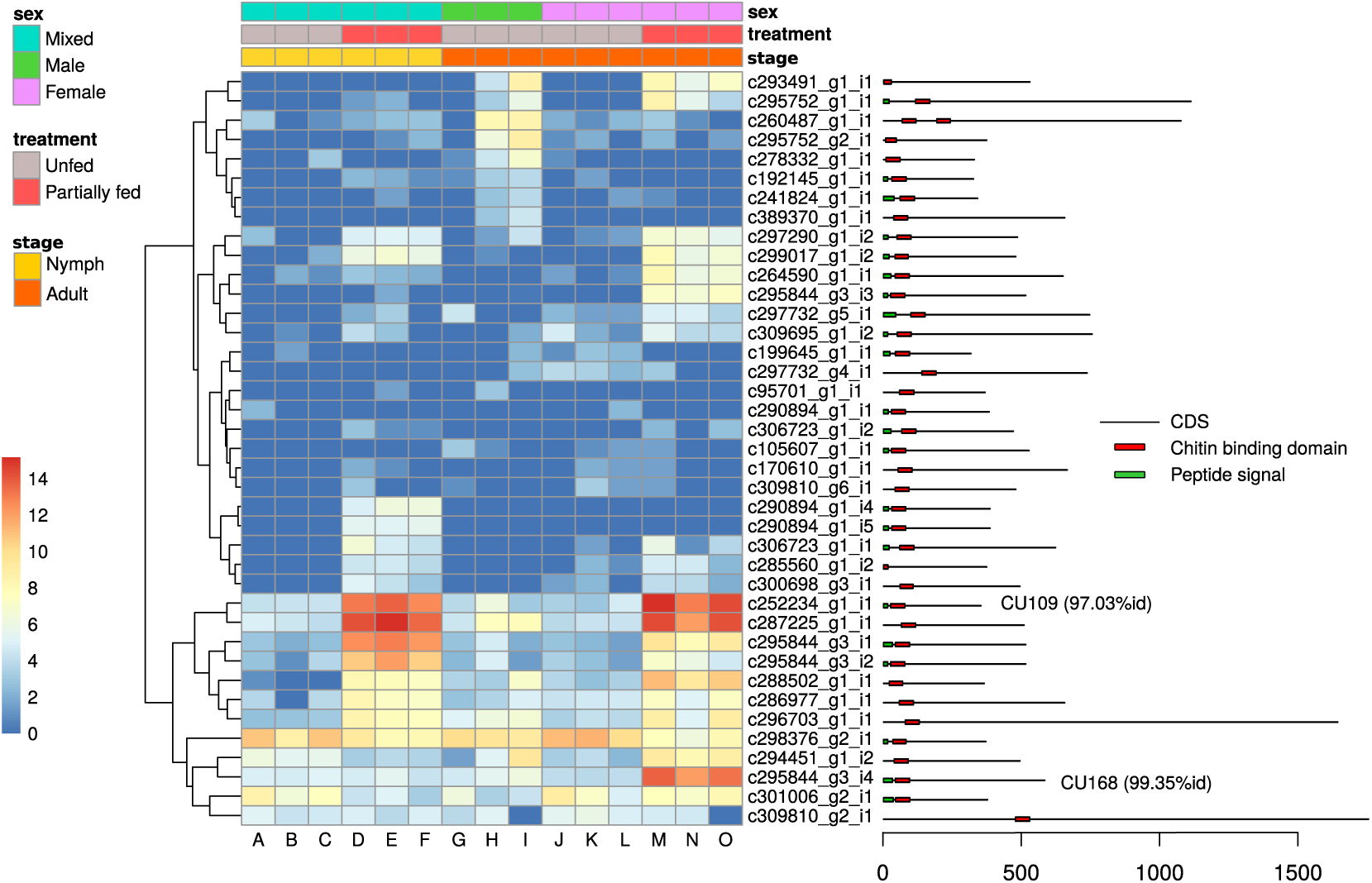
Condition-dependent expression of genes that contain a chitin-binding domain (PFAM domain PF00379). The first three rows above the heat map indicate the sex, seeding treatment and stage condition of the *I. ricinus* ticks in each of the 15 libraries (columns A to O), according to a color legend detailed on the left. Each row of the heat map corresponds to a different gene (contig). Colors of the cells represent the level of expression (log(“rax-count”+1)) of the gene in each library (counted by Kallisto). The legend on the right shows the position of the signal peptide domain and of the chitin-binding domain on the coding sequence of the transcript (CDS). Best-hit UNIPROT names are indicated for genes with more than 90% protein identity.

**Figure 5:**
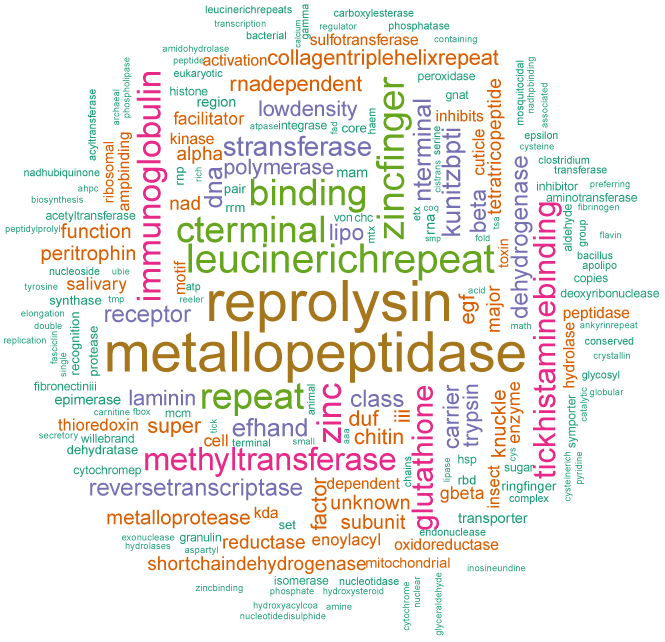
Relative abundance of terms in the identified PFAM domains of transcripts over-expressed in partially fed ticks. Transcripts over-expressed during blood feeding were defined as those transcripts that were 4-fold more abundant in partially fed ticks compared to unfed ticks and that had a p-value < 0.05 (DESeq2 analysis). Terms were extracted from PFAM annotations and the first 10 domains were retained for each transcript. Terms or characters that were not informative (e.g. “protein”, “domain”, “motif”, “dots”, etc.) were trimmed.

### Polymorphism

Starting from the 368,367 SNPs initially detected by KisSplice using the default parameters, we applied more stringent criteria to increase the specificity: only 8,955 SNPs were supported by at least 10 reads in each strain. We further filtered out all SNPs not assigned unambiguously to a single component of the T-DBG, resulting in a final set of 5,422 SNPs. For this smaller but very robust set of polymorphic sites, the minor allele frequency (MAF) and heterozygosity were computed. A factorial component analysis (FCA) showed that the three wild strains of *I. ricinus* (SEN, CZ-W, and LUX) were grouped very closely together (Figure 6). These three wild tick strains had very similar levels of heterozygosity across loci on the three planes representations (axis 1 versus axis 2 plane, axis 1 versus 3, and axis 2 versus 3 of the FCA). By contrast, the lab strains (CZ-F1, NEU, and CL) were distant from the central cluster of wild strains in the axis 1 versus axis 2 plane space, and were also distant from each other. These lab strains formed a separate group supported by the third axis (see plane space representation of corresponding to axis 1 versus 3 and axis 2 versus 3). *F_ST_* calculated by BayeScan showed very low values for the wild strains (SEN, LUX, CZ-W), but high values for the lab strains (CZ-F1, NEU, and CL) (see supplementary Figure S6). This result is in agreement with the FCA, and suggests that the three lab strains are differentiated from a typical wild population and are also different from each other. Density of the loci’s heterozygosities for each strain showed similar patterns (see supplementary Figure S7) with. Again, the three wild strains exhibiting showed similar distributions of heterozygosity whereas the two of the lab strains (CZ-F1 and NEU) showed very similar profiles: a high proportion of sites with very low heterozygosity (fixed or nearly-fixed SNPs) and a low proportion of sites with intermediate levels of heterozygosity. The CL strain showed an even more striking increase in the proportion of fixed sites. All these results showed that the three lab strains have less heterozygosity, differ strongly from wild ticks, and differ strongly from each other.

## Discussion

Several recent RNAseq studies on *I. ricinus* have focused on the transcriptomes of specific tissues such as the midgut, salivary glands, haemocytes, and ovaries [13, 14, 15, 16, 17, 18, 19]. In contrast, we tried to obtain a broader picture of the transcriptome of *I. ricinus* by sequencing and assembling transcripts from whole ticks in different conditions defined by developmental stage, sex, and feeding status. First, our *de novo* assembly showed high overall completeness and significantly enlarged the catalogue of known coding sequences for *I. ricinus*. Second, we detected blood feeding-induced changes in gene expression at the level of the whole body. Third, our analysis of polymorphism in the transcriptome sequence data of our study and previously published studies allowed us to compare the levels of genetic diversity between outbred wild strains of *I. ricinus* versus inbred lab strains and tick cell lines.

### Genomic Resources for transcribed sequences

Using the BUSCO approach, our assembly showed high completeness with metrics that compared very favorably with previously published *de novo* assemblies (TSA contigs). All of these previous assemblies were based on specific tick tissues (midgut, salivary glands, haemocytes, etc.), which probably contain a smaller repertoire of transcripts which could explain their lower level of completeness. We also suggest that the sequencing strategy used for these assemblies, a combination of 454 and Illumina technologies, may have produced sub-optimal results. For exemple the error-rich 454 sequences may have caused numerous indels in the final contigs, causing difficulties for ORF prediction [46]. In contrast, we found that 231 BUSCO genes were absent from our assembly but present in the published TSA. One explanation is that tissue-specific transcriptomes (previous studies) will detect transcripts that may be rare at the level of the whole body (our study). A second explanation is the strong time-dependence of gene expression during the blood meal as shown in previous studies on *I. ricinus* [14] and *I. scapularis* [47]. Our experimental design had only a single time point for nymphs and adults ticks, which might have resulted in missing some of the transcripts. This observation suggests that it is important to combine differences sources of data to obtain the most complete description of the transcriptome of the species. Overall, one third of the assembled transcripts showed similarity with known proteins and 70.8% of the contigs larger than 1 kb were successfully annotated. The majority of the best hits (70.0%) were matches to *Ixodes* tick species, and this result was expected given that a complete genome sequence is available for *I. scapularis* [48]. A smaller fraction had best matches to other groups, primarily arthropods (19.6%). These matches could be genes that are not fund in other tick genomic resources due to their relative incompleteness or to the true absence of homologous genes in other tick species. A small number (n=366) of contigs were assigned to genes of mice and rabbit (on which the nymphs and adults had fed), which indicates that host RNA was ingested during the blood meal. As expected, the mean proteic identity reflected phylogenetic distance, with one exception. We found a higher mean proteic identity of our transcripts to *I. scapularis* than to *I. ricinus*. One explanation for this counter-intuitive result is that the available genomic resources are much greater for *I. scapularis* than *I. ricinus*. A complete genome has been published for *I. scapularis* [48], but only a genome survey is available for *I. ricinus* [17, 49]. For *I. ricinus*, most proteic sequences in the databases are derived from *de novo* assembled transcriptomes that are still rather incomplete. Thus, one explanation for the relatively low mean identity of the hits to *I. ricinus* could be the absence of the same gene in the data banks (matches would often correspond to paralogous gene copies or shared proteic domains).

**Figure 6:**
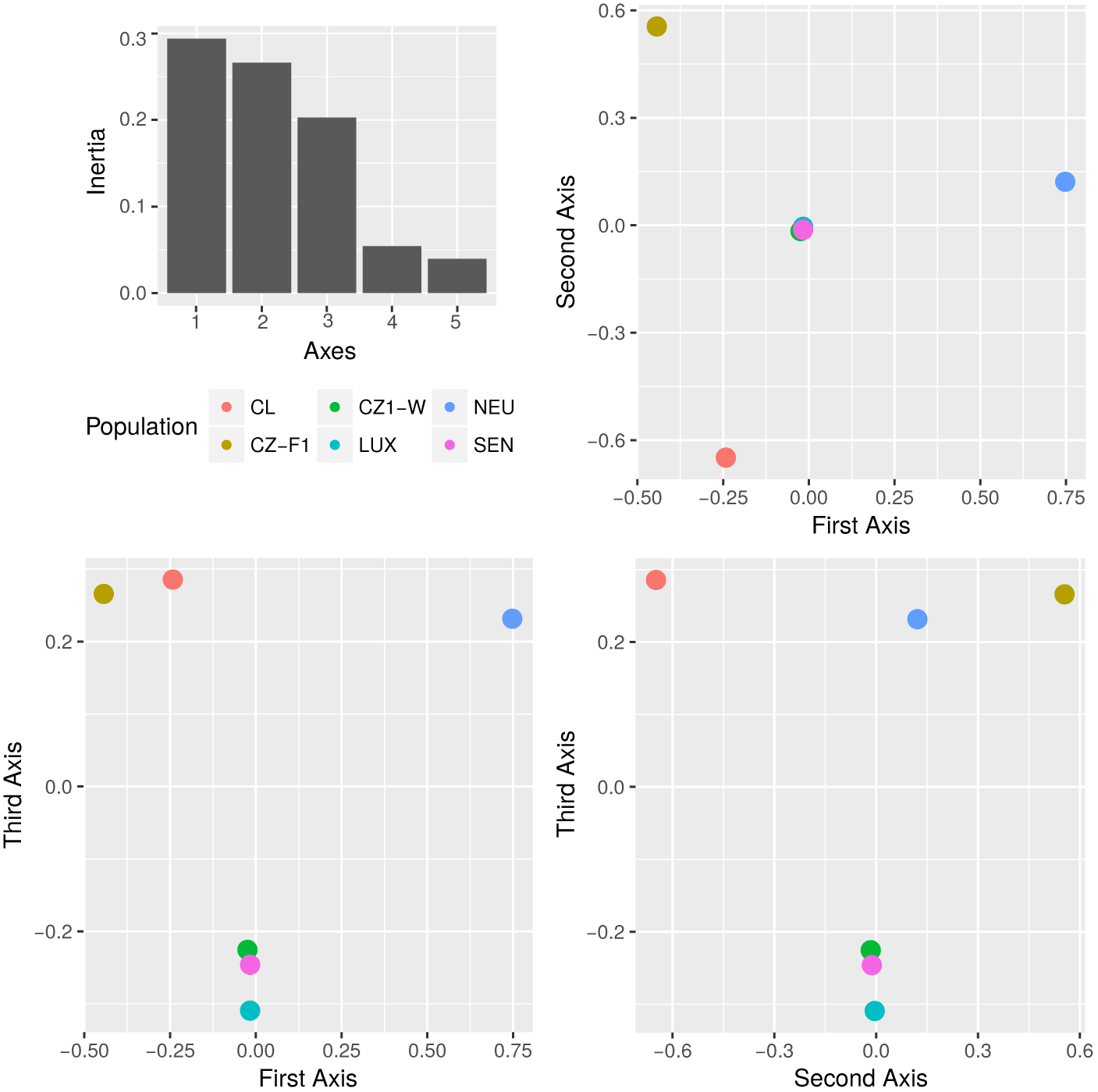
Results of the factorial component analysis (FCA). The FCA explored how six different RNAseq data sets of *I. ricinus* are grouped based on the heterozygosity at the single nucleotide polymorphism (SNP) loci. The variables correspond to heterozygosity at the 5,422 selected SNPs.

### Gene expression in response to blood feeding

Differential expression analysis showed that 11,322 transcripts were either up- or down-regulated during blood feeding. Blood feeding induced more changes in gene expression than stage (two times more) or sex (almost six times more). The enrichment tests for unfed and partially fed conditions compared the GO terms between the differentially expressed transcripts and 39,719 unbiased transcripts. The most important enriched GO terms were related to production of the tick cuticle. Over the course of the blood meal (3 to 8 days depending on the stage), hard ticks undergo dramatic changes in body size. When adult females reach repletion, their body weight increases by a factor of 100 [50]; which means that ticks must completely remodel their cuticle during repletion. Previous studies on adult female *I. ricinus* ticks observed an increase in cuticle thickness from 30 *µm* to 105 *µm* during the slow phase (first phase) of engorgement, followed by a decrease to 45 *µm* during the rapid phase (second phase) of engorgement [20, 45, 51, 50]. These studies support our finding that *I. ricinus* ticks increase their production of cuticular proteins during the blood meal, in order to greatly expand their body size.

Several studies of the transcriptome of *I. ricinus* during the blood meal have shown the prominent role of several functional groups [52], including metalloproteases [53] and proteases inhibitors, such as the Kunitz-BPTI group [54, 55]. Unexpectedly, these groups of genes were not identified in the GO-enrichment tests made for the comparison between unfed and partially fed tick ticks (if we consider only the statistically significant comparisons after correcting for multiple testing). However, examining the complete list of GO terms (for Molecular Function) associated with up-regulated transcripts in the fed condition revealed that the term GO:0004222 (metalloendopeptidase activity) appeared in fourth position, after three terms associated with cuticle production (Table S5). There are several reasons why metalloproteases and protease inhibitors did not appear in the list of genes that were significantly differentially expressed between conditions. One reason is the relatively limited fraction of genes with a GO assignation(15%), which resulted in a lack of power for these comparisons. A second reason is the fact that the previous studies investigated specific tissues (e.g. salivary glands) whereas our study concerned whole ticks. If expression of secreted BP Kunitz proteins is mostly restricted to the tick salivary glands, then any change in gene expression would be diluted when considering the whole tick body where other metabolic processes dominate, such as those related to cuticle production. A third reason is that GO terms associated with BP Kunitz proteins, metalloproteases, reprolysin, etc, were not exclusive to upregulated in the partially fed ticks. In fact, several of the genes that were considered as unbiased or genes that were strongly up-regulated in unfed ticks had the same GO terms. The same observation applies to other terms that have been associated with blood-feeding in previous studies, or to other terms (e.g. zinc finger). The text mining analysis of the PFAM domains also found a high frequency of terms like reprolysin and Kunitz domains in the up-regulated transcripts of the partially fed ticks. This result is in agreement with previous publications [54, 55, 53]. However, we note that these domains are also common in the up-regulated transcripts in unfed ticks. The text mining analysis found that some terms are abundant across the different states of developmental stage, sex, and feeding condition. For example, the term “zinc finger” was the most abundant term in three different states: nymph, adult, and female. This observation suggests that ticks contain multigenic families that share these common domains and that different genes in these families are expressed at different points in the tick life cycle. This observation also suggests a subtle transcriptional landscape, where shifts in gene expression should be studied at the level of the gene rather than at the level of blocks of genes sharing the same GO or domain assignation.

### Genetic Diversity

Our study of single nucleotide polymorphisms found substantial differences in heterozygosity between the different sources of *I. ricinus* used in RNAseq studies. The factorial component analysis indicated that three data sets corresponding to wild ticks (SEN, CZ-W, and LUX) clustered together, whereas each of the three lab strains (NEU, CZ-F1, and CL) differed markedly from that cluster and from each other. The clustering of the wild tick tick populations indicates that they have similar levels of heterozygosity (and should not be interpreted as the absence of genetic diversity between the three wild tick populations). In contrast, the lab lines show reduced levels of heterozygosity and frequent allele fixation at each of the SNP sites (Figure S7), 5 and reduced levels of heterozygosity. Reduction in heterozygosity was expected in the CZ-F1 strain because it was derived from a single mating of two wild ticks. Reduced heterozygosity was also expected in the NEU strain because it was derived from a laboratory colony that has a long history of inbreeding and small effective population size. The material from the tick cell line (CL) had an even more extreme reduction of heterozygosity, which is an feature of cell lines [56]. The lab strains differ from each other because of genetic drift, which results in the random fixation of alleles in each lab strain. Our study illustrates that sequencing the transcriptome of tick populations allows the genotyping of thousands SNPs at hundreds of genes, which can greatly extend the traditional populations genetic approaches based on much fewer genetic markers [14, 43, 57].

## Conclusion

Our *de novo* assembly showed high overall completeness and significantly enlarged the catalogue of known coding sequences for *I. ricinus*. Our study investigated the transcriptome of whole ticks that differed with respect to their developmental stage, sex, and feeding condition. This approach allowed us to detect changes in gene expression at the level of the whole body instead of specific tissues. We found that blood feeding induced a strong up-regulation of transcripts associated with cuticle production. Finally, our analysis of polymorphism in the transcriptome sequence data of our study and previously published studies allowed us >5,000 robust SNPs from coding regions, and to compare the levels of genetic diversity between outbred wild strains of *I. ricinus* versus inbred lab strains and tick cell lines.

## Availability of data and materials

Reads from sequencing the 15 libraries were deposited in the SRA section of NCBI under the BioProject ID: PRJNA395009. The contigs are available under accession GFVZ00000000 (NCBI, TSA). The annotation table was deposited on the Zenodo repository with a DOI (10.5281/zenodo.1137702), available on https://zenodo.org/record/1137702.

## Authors’ contributions

CR and OP conceived and designed the experiments; OR and MJV reared the ticks and provided the raw material for this study; AD and CH performed the molecular work (RNA extraction). NPC, MC and CR analyzed the data. NPC and CR wrote the paper. All authors read and revised the manuscript and approved its final version.

## Acknowledgements

We are grateful to the Genotoul bioinformatics platform Toulouse Midi-Pyrénées (Bioinfo Genotoul) for providing support and computing resources. Part of the computations (multiple transcriptome assembly for subsamples of reads) were performed at the Bordeaux Bioinformatics Center (CBiB).

